# Analysis of Ultrasonic Vocalizations from Mice Using Computer Vision and Machine Learning

**DOI:** 10.1101/2020.05.20.105023

**Authors:** Antonio H. O. Fonseca, Gustavo M. Santana, Sérgio Bampi, Marcelo O. Dietrich

## Abstract

Mice emit ultrasonic vocalizations (USV) to transmit socially-relevant information. To detect and classify these USVs, here we describe the development of VocalMat. VocalMat is a software that uses image-processing and differential geometry approaches to detect USVs in audio files, eliminating the need for user-defined parameter tuning. VocalMat also uses computational vision and machine learning methods to classify USVs into distinct categories. In a dataset of >4,000 USVs emitted by mice, VocalMat detected more than >98% of the USVs and accurately classified ≈86% of USVs when considering the most likely label out of 11 different USV types. We then used Diffusion Maps and Manifold Alignment to analyze the probability distribution of USV classification among different experimental groups, providing a robust method to quantify and qualify the vocal repertoire of mice. Thus, VocalMat allows accurate and highly quantitative analysis of USVs, opening the opportunity for detailed and high-throughput analysis of this behavior.

## 1 Introduction

Vertebrates use vocal communication to transmit information about the state of the caller and influence the state of the listener. This information can be relevant for the identification of individuals or groups [18]; status within the group (e.g.: dominance, submissive, fear or aggression) [25]; next likely behavior (e.g.: approach, flee, play or mount) [22]; environment conditions (e.g.: presence of predators, location of food) [33]; and facilitation of mother-offspring interactions [12].

Mice emit ultrasonic vocalizations (USVs) in a frequency range (≈30 – 110kHz) above the human hearing range (≈2 – 20kHz) [39, 23, 25, 26, 24, 29, 5, 17, 13, 6]. These USVs are organized in *phrases* or *bouts* composed by sequences of *syllables*. The syllables are defined as continuous units of vocal sound not interrupted by a period of silence. The syllables are composed of one or more notes and are separated by salient pauses and occur as part of sequences [3, 19]. These transitions across syllables do not occur randomly [19, 8], and the changes in the syllables sequences, prevalence and acoustic structure match current behavior [9], genetic strain [36, 30], and developmental stage [16]. USVs are most commonly emitted by mouse pups [31] and are modulated during development [16, 14, 8]. In the adult mouse, USVs are emitted in both positive and negative contexts [2]. Thus, understanding the complex structure of USVs emitted by mice is key to advancing vocal and social communication research in mammals.

In the past years, tools for USV analysis advanced significantly [10, 36, 22, 9, 3, 34]. In terms of USV detection, the majority of the software tools available depend on user inputs [22, 36, 34] or present limited detection capabilities [3, 9]. An exception is DeepSqueak [10], which uses automated detection of USVs from audio recordings. Regarding USV classification, no consensus exists on the biological function of the various USV sub-classes, making it challenging to develop a tool for all purposes. Thus, different tools use supervised [3, 9, 10] and unsupervised [7, 36, 10] methods to classify USVs into different syllable classes. Our goal was to create a tool with high accuracy for USV detection that allows for the flexible use of any classification method.

Here, we describe the development of VocalMat, a software for robust and automated detection and classification of mouse USVs from audio recordings. VocalMat uses image-processing and differential geometry approaches to detect USVs in spectrograms, eliminating the need for parameter tuning. VocalMat shows high accuracy in detecting USVs, outperforming previous tools. This high accuracy allows the use of multiple tools for vocal classification. In the current version of VocalMat, we embedded a supervised classification method that uses computer vision techniques and machine learning to label each USV in eleven different sub-classes. The output of the vocal classification provides the additional benefit of a probability distribution of vocal classes, allowing for the use of nonlinear dimensionality reduction techniques to analyze the vocal repertoire. We provide an example of such analysis by applying Diffusion Maps and Manifold Alignment to an experimental dataset. Thus, VocalMat is a highly accurate software to detect and classify mouse USVs in an automated and flexible manner.

## 2 Results

### 2.1 Detection of mouse USVs using imaging processing

VocalMat uses multiple steps to analyze USVs from vocalizing mice in audio files (see Figure 1A for the general workflow). Initially, the audio recordings are converted into high-resolution spectrograms through a short-time Fourier transformation (see Methods and Materials). The resulting spectrogram consists of a matrix, wherein each element corresponds to an intensity value (power spectrum represented in decibels) for each time-frequency component. The spectrogram is then analyzed in terms of its time-frequency plane, where high-intensity values are represented by brighter pixels in a gray-scale image (Figure 1B). The gray-scale image undergoes contrast enhancement and adaptive thresholding for binarization (see Methods and materials). The segmented objects are further refined via morphological operations (Figure 1C and Figure S1), thus resulting in a list of segmented blobs (hereafter referred to as USV candidates) with their corresponding spectral features (Figure 1D).

**Figure 1:**
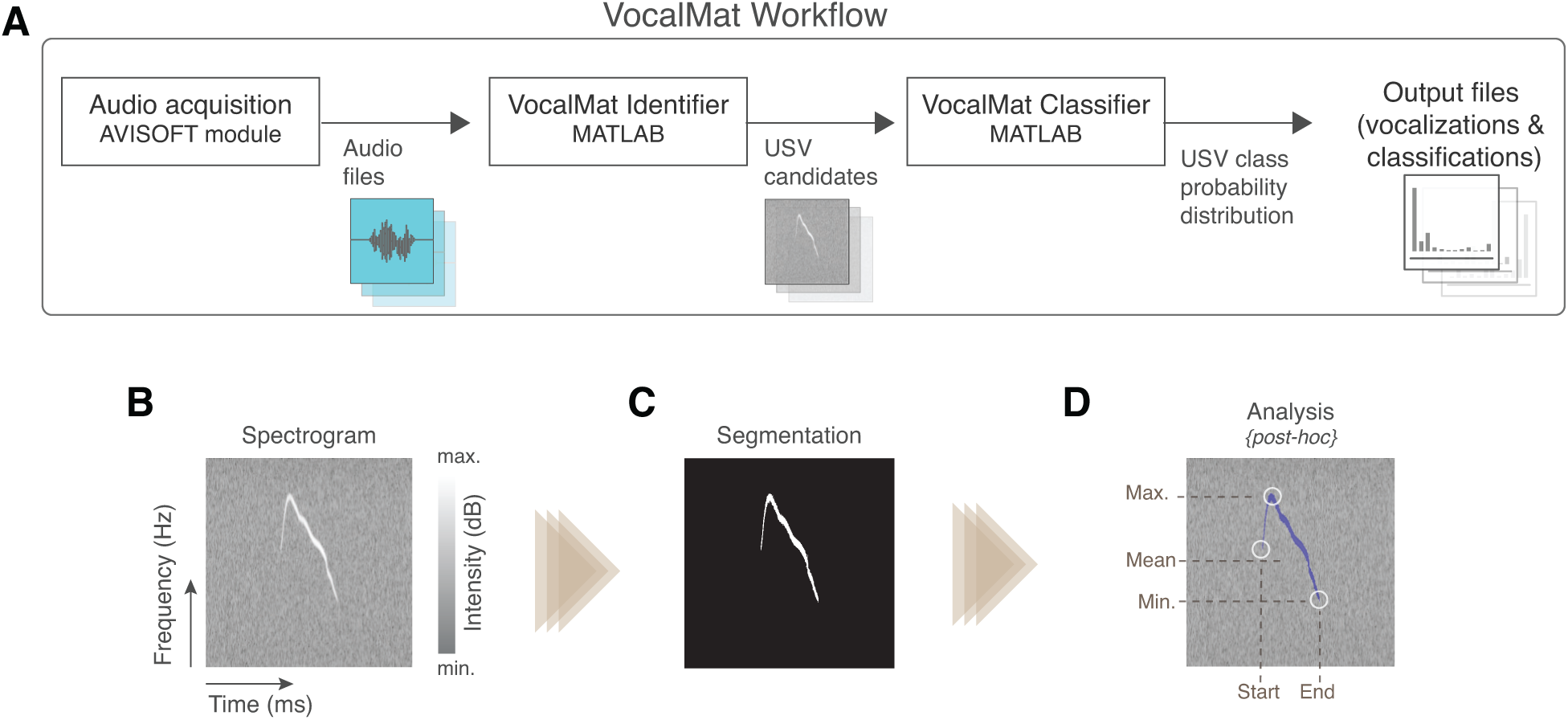
Overview of the VocalMat pipeline for USV detection and analysis. **(A)** Workflow of the main steps used by VocalMat, from audio acquisition to data analysis. **(B)** Illustration of a segment of spectrogram. The time-frequency plan is depicted as a gray scale image wherein the pixel values correspond to intensity in decibels. **(C)** Example of segmented USV after contrast enhancement, adaptive thresholding and morphological operations (see Figure S1 for further details of the segmentation process). **(D)** Illustration of some of the spectral information obtained from the segmentation. Information on intensity is kept for each time-frequency point along the segmented USV candidate.

This list of USV candidates may contain noise (i.e., detected particles that are not part of any USV) and multiple candidates that correspond to the same USV. To address this, a minimum of 10 ms interval between two successive and distinct syllables is assumed based on experimental observations [9]. To reduce the amount of data stored for each USV, the features extracted from detected candidates are represented by a mean frequency and intensity every 0.5 ms. The means are calculated for all the individual candidates, including the ones overlapping in time, hence preserving relevant features such as duration, frequency, intensity, and harmonic components (Figure 1D).

Harmonic components are also referred to as nonlinear components or composite [31, 30]. Here, we did not consider harmonic components as a different syllable, but rather as an extra feature of a syllable [16]. Therefore, each detected USV candidate may or may not present a harmonic component. A harmonic component was considered as a continuous USV candidate (i.e., no discontinuities in time and/or frequency) overlapping in time with the main component of the USV (similar to [16]).

Besides the list of USV candidates and their spectral features, the segmentation process also exports image files of 227 x 227 pixels, in which the USV candidate is centralized in windows of 220 ms (see Figure 1B). This temporal length is defined as twice the maximum duration of USVs observed in mice [16], thus preventing cropping.

### 2.2 Eliminating noise via Local Median Filter

Initially, we used VocalMat to detect USVs in a set of 64 recordings, resulting in a pool of 59,781 USV candidates, which includes real USVs and noise (Figure 2A and Methods and Materials). Visual inspection of the dataset revealed that artifacts generated during the segmentation process dominated the pool of USV candidates (see Figure 2B for examples of real USVs and noise in the pool of USV candidates). This type of artifact is characterized by its low intensity compared to real USVs. To remove these artifacts from the pool of USV candidates, we applied a ‘Local Median Filter’ step, a method to estimate the minimum expected contrast between a USV and its background for each audio recording. This contrast is calculated based on the median intensity of the pixels in each detected USV candidate *k* (referred to as 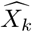), and the median intensity of the background pixels in a bounding box containing the candidate *k* (referred to as 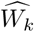) (Figure 2C). Thus, the contrast is defined as the ratio 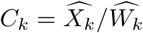.

**Figure 2:**
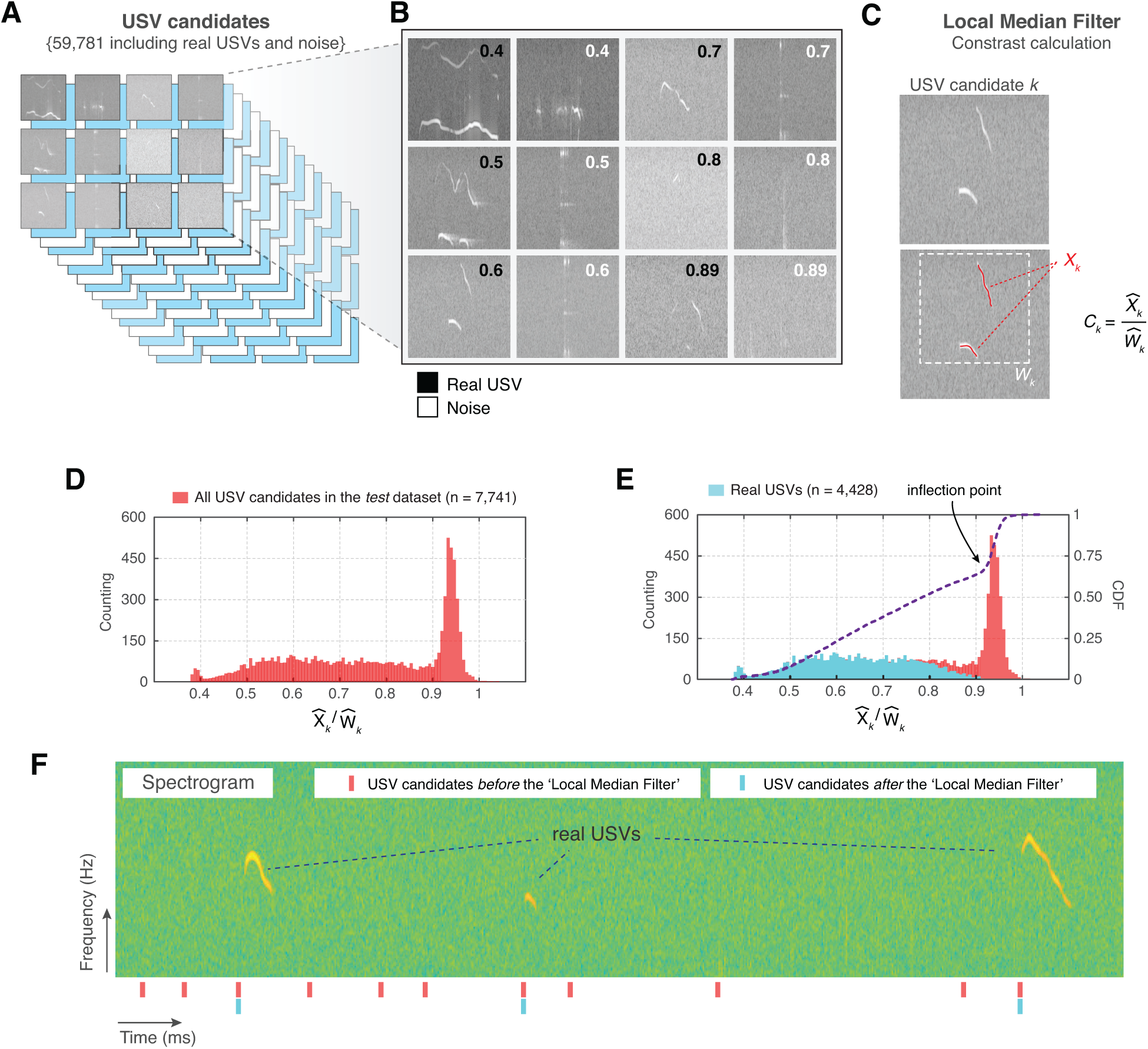
Noise elimination process for USV candidates. **(A)** In a set of 64 audio files, VocalMat identified 59,781 USV candidates. **(B)** Examples of USVs among the pool of candidates that were manually labeled as either noise or real USVs. The score (upper-right corner) indicates the calculated contrast *C*_*k*_ for the candidate. **(C)** Example of contrast calculation (*C*_*k*_) for a given USV candidate *k*. The red dots indicate the points detected as part of the USV candidate (*X*_*k*_) and the dashed-white rectangle indicates its evaluated neighborhood (*W*_*k*_). **(D)** Distribution of the *C*_*k*_ for the USV candidates in the *test* dataset. **(E)** Each USV candidate was manually labeled as real USV or noise. The distribution of *C*_*k*_ for the real USVs (cyan) compared to the the distribution for all the USV candidates (red) in the *test* dataset. The blue line indicates the cumulative distribution function (CDF) of *C*_*k*_ for all the USV candidates. The inflection point of the CDF curve is indicated by the arrow. **(F)** Example of a segment of spectrogram with 3 USVs. The analysis of this segment without the ‘Local Median Filter’ results in an elevated number of false positives (noise detected as USV). ‘Red’ and ‘cyan’ ticks denote the time stamp of the identified USV candidates without and with the ‘Local Median Filter’, respectively.

To validate this method, we manually inspected the spectrograms and labeled USV candidates in a subset of audio files (hereafter referred to as *test* dataset and described in Table 1). A total of 7,741 USV candidates were detected using the segmentation process described above, representing 1.75 times more USV candidates than the manual counting (4,441 USVs). Importantly, the segmentation step included 4,428 real USVs within the pool of USV candidates, therefore missing 13 USVs. Thus, the segmentation process of VocalMat presents a rate of missing USVs of 0.29%.

**Table 1:** Summary of experimental conditions covered in the test dataset

| Age | Microphone gain | Chamber | Heating |
| --- | --- | --- | --- |
| P9 | Maximum | Yes | No |
| P9 | Maximum | Yes | No |
| P9 | Maximum | Yes | No |
| P10 | Intermediary | No | No |
| P10 | Intermediary | No | No |
| P10 | Maximum | Yes | Yes |
| P10 | Maximum | Yes | Yes |

The distribution of *C*_*k*_ for real USVs and for noise showed that the peak at high *C*_*k*_ (i.e., low contrast) in the distribution was dominated by USV candidates corresponding to artifacts of the segmentation process (Figure 2D-E). The *C*_*k*_ of real USVs (mean = 0.642, SEM = 1.841 *×* 10^−3^, median = 0.640, 95% CI [0.638, 0.646]; N = 4,428) was significantly lower than the *C*_*k*_ of noise (mean = 0.922, SEM = 9.605 *×* 10^−4^, median = 0.936, 95% CI [0.921, 0.924]; n = 3,336; *P* < 10^−15^, *D* = 0.894, Kolmogorov-Smirnov test; Figure 2D-E). This unbalanced bimodal distribution causes an inflection point on the cumulative distribution function (CDF) of *C*_*k*_ that matches the ratio observed for segmentation artifacts (Figure 2E). Therefore, based on these results, we used the calculated inflection point as a threshold to effectively eliminate a substantial amount of noise from the pool of USV candidates (details on this calculation are provided in Methods and Materials).

In the *test* dataset, 5,171 out of 7,741 USV candidates survived the ‘Local Median Filter’. This number includes real USVs (4,421) and remaining noise of lower *C*_*k*_. This remaining noise presented high intensity and commonly originated from external sources (Figure 2B, E). The 7 real USVs eliminated in this step presented a high *C*_*k*_ (mean = 0.942, SEM = 5.871 *×* 10^−3^, median = 0.943, 95% CI [0.927, 0.956]; n = 7). To illustrate the performance of the ‘Local Median Filter’, Figure 2F shows a segment of a spectrogram with 3 real USVs and a total of 11 USV candidates detected. After applying the ‘Local Median Filter’, only the real USVs remained in the pool of USV candidates. Thus, the ‘Local Median Filter’ effectively eliminates segmentation noise from the pool of USV candidates, which provides two main advantages: it decreases the number of USV candidates used in downstream analysis and, consequently, reduces the number of false positives.

In an ideal experimental setting with complete sound insulation and without the generation of noise by the movement of the animal, no further step is required to identify real USVs using VocalMat. Since this is difficult in experimental conditions, we applied a second step in the noise elimination process.

### 2.3 Using Convolutional Neural Network for noise identification

To identify USVs in the pool of USV candidates that passed the ‘Local Median Filtering’, we trained a Convolutional Neural Network (CNN) to classify each USV candidate into one of 11 USV categories or noise (see Figure 4A for examples of the different USV categories). We used a dataset containing 10,871 samples manually labeled as one of the 11 USV categories and 2,083 samples of noise (see Methods and materials). The output of the CNN is the probability of each USV candidate belonging to one of the 12 categories. The most likely category defines the label of the USV candidate (Figure 3A).

**Figure 3:**
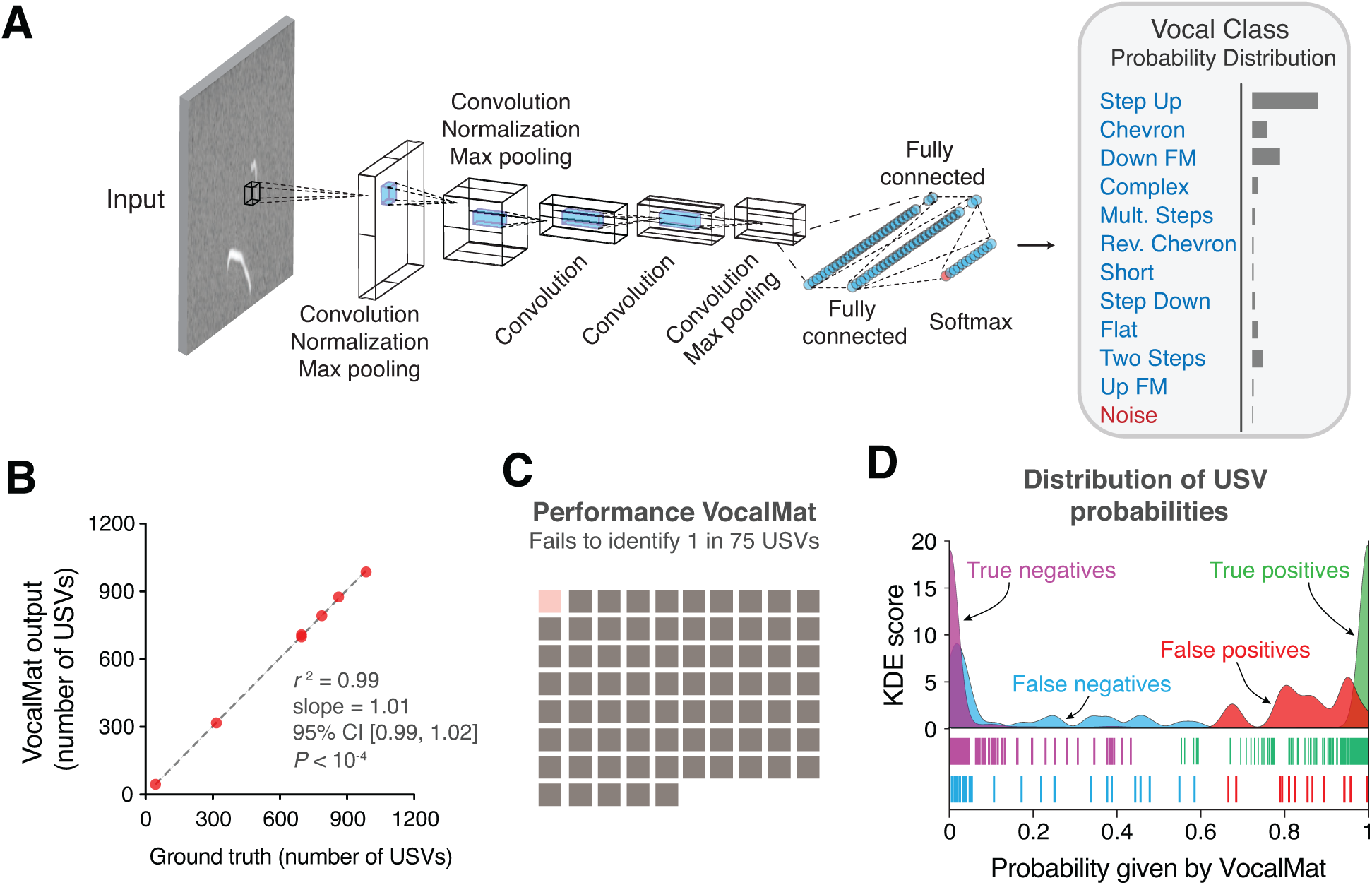
VocalMat USV classification using a Convolutional Neural Network. **(A)** Illustration of the AlexNet architecture post end-to-end training on our *training* dataset. The last threes layers of the network were replaced in order to perform a twelve-categories (11 USV types plus noise) classification task. The output of the CNN is a probability distribution over the labels for each input image. **(B)** Linear regression between the number of USVs manually detected versus the number reported by VocalMat for the audio files in our *test* dataset. **(C)** Overall accuracy of VocalMat in detection and classification of detected USV candidates. VocalMat fails to identify 1 in every 75 USVs. **(D)** Distribution of probabilities *P* (*USV*) for the true positive (green), false positive (red), false negative (cyan) and true negative (magenta). Ticks represent individual USV candidates.

**Figure 4:**
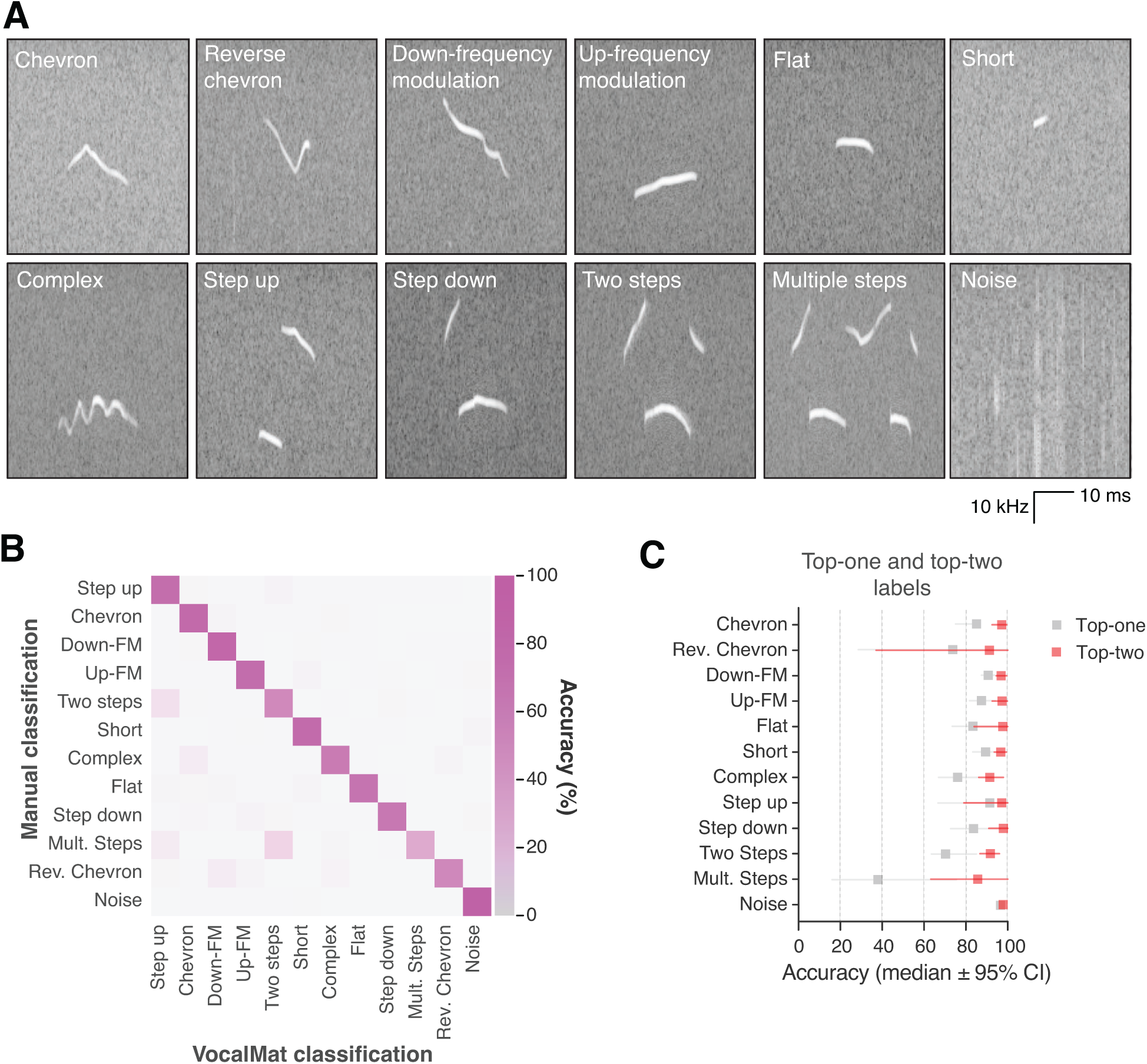
VocalMat performance for USV classification. **(A)** Example of the 11 categories of USVs plus noise that VocalMat used to classify the USV candidates. **(B)** Confusion matrix illustrating VocalMat’s performance in multiclass classification (see also Table S5). **(C)** Comparison of classification performance for labels assigned based on the most likely label (Top-one) versus the two most likely labels (Top-two) (see Table S6). Symbols represent median ±95 % confidence intervals.

To evaluate the performance of VocalMat in distinguishing between USVs and noise, we used the 5,171 USV candidates in the *test* dataset that passed the ‘Local Median Filter’ step (Methods and materials). For the detection evaluation, we compared the score for the label ‘Noise’ (*P* (*Noise*)) to the sum over the 11 USV categories (*P* (*USV*)). The rate of detected USVs labeled as such (true positive or sensitivity) was 99.04 ± 0.31% (mean ± SEM; median = 99.37; 95% CI [98.27, 99.80]). A linear regression analysis between manually validated data from different audio files and the true positives of the CNN revealed an almost-perfect linearity (*r*^2^ = 0.99, 95% CI [0.99, 1.02]), *P* < 10^−4^, and slope *α* = 1.01), suggesting high accuracy of VocalMat in detecting USVs from audio files and removing noise (Figure 3B). The rate of detected USVs labeled as noise (false negative) was 0.96 ± 0.31% (mean ±SEM; median = 0.61; 95% CI [0.20, 1.73]). The rate of detected noise labeled as noise (true negative rate or specificity) was 94.40 ± 1.37 % (mean ± SEM; median = 95.60; 95% CI [91.60, 97.74]). The rate of detected noise labeled as USV (false positive) was 5.60 ± 1.37% (mean ± SEM; median = 4.40; 95% CI [2.26, 8.94]), representing a total of 42 wrongly detected USVs out of the 5,171 USV candidates in the *test* dataset. Finally, the rate of USVs not detected (missed rate) was 0.28 ± 0.09% (mean ± SEM; median = 0.23; 95% CI [0.05, 0.51]). All together, the overall accuracy in identifying USVs was 98.63 ± 0.20% (mean ± SEM; median = 98.55; 95% CI [98.14, 99.11]) for manually validated audio files. Thus, VocalMat fails to identify approximately 1 in 75 USVs (Figure 3C).

### 2.4 Characteristics of mislabeled USV candidates by VocalMat

We further calculated other measures of performance (Figure 3D). For USVs wrongly labeled as noise (false negative), the probability of being a USV was 0.15 ± 0.03 (mean ± SEM; median = 0.04; 95% CI [0.09, 0.22]; Figure 3D), while for noise labeled as USV (false positive), the probability of being USV was 0.85 ± 0.03 (mean ± SEM; median = 0.86; 95% CI [0.80, 0.91]; Figure 3D). These probabilities contrast with cases in which VocalMat correctly identified USV and noise. USVs that were correctly identified had a probability of being USV of 0.99 ± 3.78 × 10^−4^ (mean ± SEM; median = 1.00; 95% CI [0.99, 0.99]; Figure 3D). Noise that was correctly identified had a probability of being noise of 0.99 ± 1.78 × 10^−3^ (mean ± SEM; median = 1.00; 95% CI [0.98, 0.99]; Figure 3D). These results indicate that the probability assigned by VocalMat flags likely errors in classification. These *flagged* candidates (i.e., with assigned low probability) can be manually inspected to correct the misclassification and retrain VocalMat.

### 2.5 Performance of VocalMat compared to other tools

In order to evaluate the performance of VocalMat in detecting USVs compared to other published tools, we analyzed the same *test* dataset with Ax [22], MUPET [36], USVSEG [34], and DeepSqueak [10]. We adopted the same validation criterion used for VocalMat (see Methods and materials).

Ax requires a series of manual inputs for their detection algorithm [22]. We tried three different settings to maximize the number of detected USVs compared to the ground-truth (Table S1). Combining the best configurations tested, the percentage of missed USVs was 4.99 ± 1.34 % (mean ± SEM; median = 4.07, 95% CI [1.73, 8.26]) and the false discovery rate was 37.67 ± 5.59 % (mean ± SEM; median = 42.56, 95% CI [23.99, 51.34]). Since Ax does not separate the selected USV candidates in real USV or noise, no false negative rate was calculated. In comparison to Ax, MUPET has a lower number of parameters to be set by the user. We tested eight different configurations of MUPET to measure its performance in detecting USVs in the validated *test* dataset (Table S2). Combining the best configurations tested, the percentage of missed USVs was 33.74 ± 3.81 % (mean ± SEM; median = 33.13, 95% CI [24.41, 43.07]), and false discovery rate of 38.78 ± 6.56 % (mean ± SEM; median = 32.97, 95% CI [22.72, 54.84]). Similarly to the other tools tested, USVSEG requires setting parameters manually for USV detection (Table S3). USVSEG displayed the best performance out of the manually configured tools, presenting a missed vocalization rate of 6.53 ± 2.56% (mean ± SEM; median = 4.26, 95% CI [0.26, 12.80]), and a false discovery rate of 7.58 ± 4.31% (mean ± SEM; median = 3.27, 95% CI [-2.97, 18.15]). It is important to emphasize that the tests with Ax, MUPET, and USVSEG did not explore all possible combinations of parameters, implying that better settings could potentially optimize the detection performance for our *test* dataset.

DeepSqueak does not demand a manual setting of parameters for USV detection and, similarly to VocalMat, it also relies on deep learning algorithms to detect and analyze USVs [10]. To measure the performance of DeepSqueak to detect USVs and compare to VocalMat, we correlated USVs detected by DeepSqueak with the time stamp of the USVs in our *test* dataset. Because DeepSqueak is not formally trained to identify the start time of USVs with precision, we used increasing tolerance for mismatches in the starting time (±5, ±10, ±15 and ±20 ms). Using 5 ms mismatch, the rate of missed USVs by DeepSqueak was 41.14 ± 6.30% (mean ± SEM; median = 35.99, 95% CI [25.71, 56.56]) and the rate of false discovery was 25.71 ± 6.74% (mean ± SEM; median = 20.89, 95% CI [9.20, 42.22]). With increasing tolerance (±10, ±15 and ±20 ms), we observed a gradual decrease in the rate of missed USVs and the rate of false discovery. The best values obtained were a rate of missed USVs of 27.13 ± 3.78% (mean ± SEM; median = 24.22, 95% CI [17.86, 36.40]) and a rate of false discovery of 7.61 ± 2.35% (mean ± SEM; median = 4.73, 95% CI [1.84, 13.39]). The manual inspection of the USVs detected by DeepSqueak revealed cases of more than one USV being counted as a single USV, which could lead to an inflated number of missed USVs. Since we did not train DeepSqueak with our dataset, it is possible that DeepSqueak could present better performance than what we report here when custom-trained.

To more directly compare DeepSqueak and VocalMat, we evaluated the performance of both tools on the sample audio provided by DeepSqueak [10]. First, we manually inspected the spectrogram of the sample audio and labeled the starting time of each of the 762 USVs identified. Of these 762 USVs, VocalMat detected 747 with a true positive rate of 91.73%, whereas DeepSqueak detected 608, with a true positive rate of 77.95%. Thus, these comparisons suggest that VocalMat shows an overall better sensitivity for USV detection when compared to DeepSqueak.

### 2.6 Detection of harmonic components

To measure the performance of VocalMat for detection of harmonic components, we compared the output of VocalMat with the *test* dataset. The rate of true positives was 93.32 ± 1.96 % (mean ± SEM; median = 92.18; 95% CI [88.54, 98.11]). The rate of USVs wrongly labeled as having a harmonic component (false positive) was 5.39 ± 1.18 % (mean ± SEM; median = 5.17; 95% CI [2.50, 8.27]). The rate of missed harmonic components (false negative) was 6.68 ± 1.96 % (mean ± SEM; median = 7.82, 95% CI [1.89, 11.46]). All combined, the error rate in identifying harmonic components was 12.19 ± 3.44 % (mean ± SEM; median = 11.92, 95% CI [3.34, 21.03]). Thus, VocalMat presents satisfactory performance in detecting the harmonic components of the USVs.

### 2.7 Classification of USVs in categories

To evaluate the performance of VocalMat in classifying the detected USVs in distinct categories, we compared the most likely label (Top-one) assigned by the CNN to the labels assigned by the investigators (i.e., ground-truth). The overall accuracy of the VocalMat classifier module is 86.05 % (Figure 4B-C and Table S5). VocalMat shows lower accuracy to detect rare USV types (e.g., reverse chevron; Figure 4A-C) or USVs with multiple components (e.g., multiple steps and two steps; Figure 4A-C). When we expanded our analysis to consider the two most likely labels assigned by the CNN (Top-two), the overall accuracy of VocalMat was 94.34 % (Figure 4E and Table S6). These observations suggest a possible overlap between the definition of categories. Based on these analyses, we reasoned that the distribution of probabilities for each of the 11 categories of USV types calculated by the CNN could provide a more fluid classification method to analyze the vocal repertoire of mice.

### 2.8 Using VocalMat to analyze and visualize the vocal repertoire of mice

To illustrate the use of the probability distribution of USV classification by VocalMat, we used data previously published by our group with over 45,000 USVs [38]. In this published dataset, two groups of ten days old mice were studied. At this age, mice vocalize in the ultrasonic range when separated from the nest. Two groups of mice were analyzed (control versus treatment) during two contiguous time points (baseline versus test). The difference between the two groups was that in the treatment group, a specific population of neurons in the brain was activated to induce higher rates of USV emission [38].

To visualize the probability distribution of USV classification by VocalMat, we used Diffusion Maps (see Methods and materials). Diffusion Maps is a dimensionality reduction algorithm that allows the projection of the probability distribution into a Euclidean space [11]. We compared all four experimental conditions against each other and visually verified that the manifolds representing the USV repertoires showed a degree of similarity (Figure 5A).

**Figure 5:**
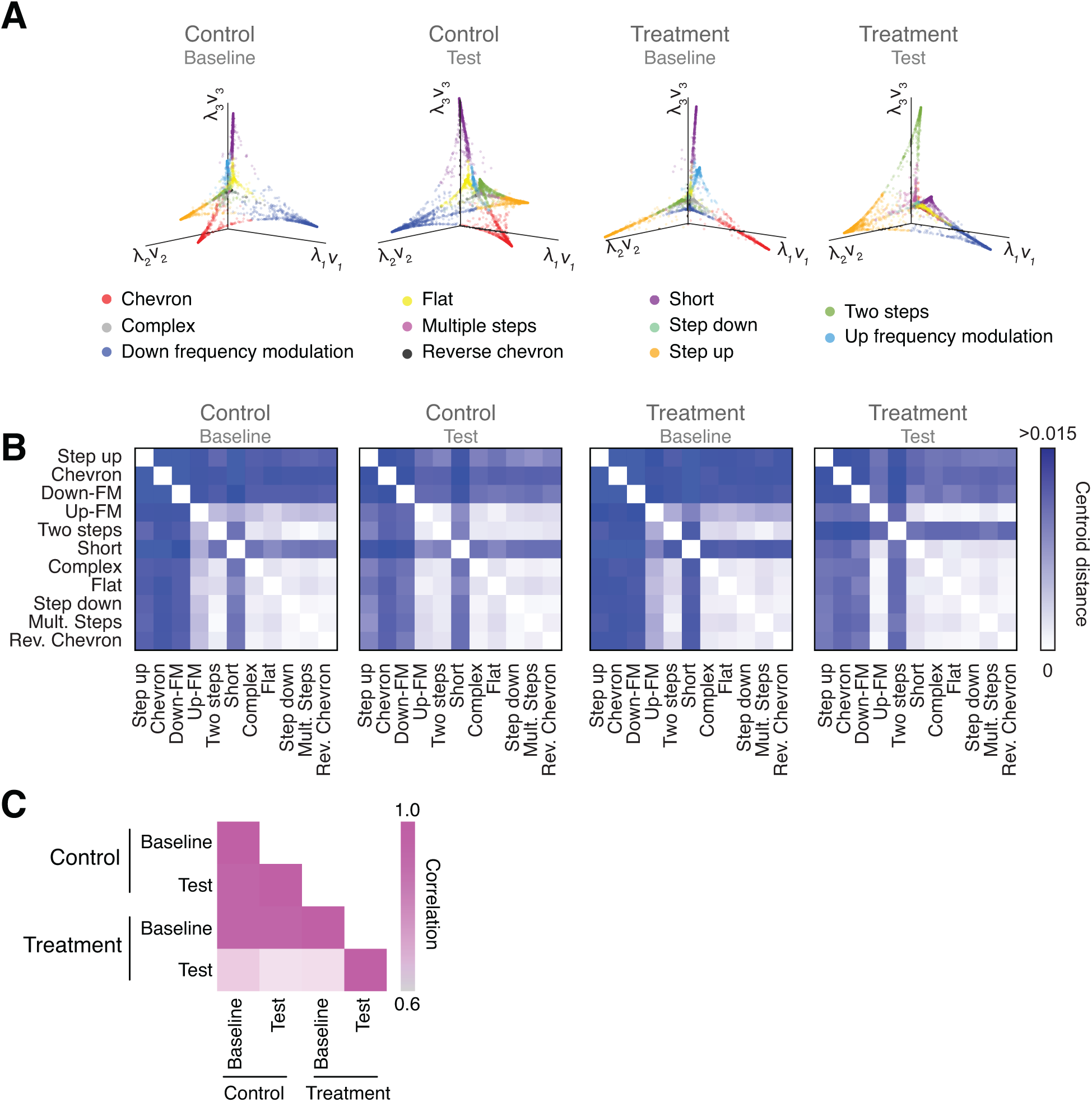
Vocal repertoire visualization using Diffusion Maps. **(A)** Illustration of the embedding of the USVs for each experimental condition. The probability distribution of all the USVs in each experimental condition is embedded in a euclidean space given by the eigenvectors computed through Diffusion Maps. Colors identify the different USV types. **(B)** Pairwise distance matrix between the centroids of USV types within each manifold obtained for the four experimental conditions. **(C)** Comparison between the pairwise distance matrices in the four experimental conditions by Pearson’s correlation coefficient.

To quantify the similarities (or differences) between the manifolds, we calculated the pairwise distance between the centroids of USV types within each manifold (Figure 5B). The pairwise distance matrices provide a metric for the manifold structure, allowing a direct comparison between the vocal repertoire of different groups. When we compared the similarity between the pairwise distance matrices in the four experimental conditions, we observed that the treatment group in the test condition presented a robust structural change in the vocal repertoire, which can be effectively represented by a matrix correlation (Figure 5C). The degree of similarity between the experimental conditions can also be visualized by comparing the structure of the manifolds. Since the manifolds are calculated separately, their coordinate system needs to be aligned to allow visual comparisons, which we achieve using the Kernel Alignment algorithm (Figure S2 and Methods and Materials) [35, 37]. The quality of the manifold alignment is assessed by

Cohen’s coefficient and overall projection accuracy into a joint space (Figure S2), showing the lowest scores for the treatment group in the test condition when compared to the other experimental conditions. Hence, these later analyses illustrate the use of the probability distribution for vocal classification and the power of dimensionality reduction techniques–such as Diffusion Maps–to provide a detailed analysis of the vocal repertoire of mice.

## 3 Discussion

We reported the development of VocalMat, a software to automatically detect and classify mouse USVs with high sensitivity. VocalMat eliminates noise from the pool of USV candidates, preserves the main statistical components for the detected USVs, and identifies harmonic components. Additionally, VocalMat’s architecture uses machine learning algorithms to classify USV candidates into 11 different USV classes or noise. VocalMat is open-source, and is compatible with high-performance computing clusters that use the Slurm job scheduler, allowing parallelized and high-throughput analysis.

VocalMat adds to the repertoire of tools developed to study mouse USVs [36, 7, 9, 3, 19, 10, 34]. We found only one study that reported the sensitivity to detect vocalizations [19]. In this manuscript, the authors reported a sensitivity of >95% compared to >98% achieved by VocalMat. Because these previous tools depend on several parameters defined by the user, it is difficult to compare their performance to VocalMat effectively. Still, our tests show VocalMat outperforming the other tools in both sensitivity and accuracy in detecting USVs without the need for parameter tuning. Moreover, VocalMat provides a flexible classification method by treating USV classification as a problem of probability distribution across different USV categories. This approach allows the analysis, visualization, and comparison of the repertoires of USVs of different mice and experimental groups using dimensionality reduction algorithms.

VocalMat uses a pattern recognition approach based on CNNs, which learns directly from the training set without the need for feature extraction via segmentation processes [32, 20]. This characteristic provides the possibility for unique adaptability of VocalMat to different experimental settings, including its use with other species and vocal types.

In summary, VocalMat is a new tool to detect and classify mouse USVs with superior sensitivity and accuracy while keeping all the relevant spectral features, including harmonic components.

## 4 Methods and Material

### 4.1 Animals

All mice used to record the emission of USV were 5-15 days old from both sexes. Dams used were 2–6 months old and were bred in our laboratory. The following mouse lines purchased from The Jackson Laboratories were used: C57Bl6/J, NZO/HlLtJ, 129S1/SvImJ, NOD/ShiLtJ, and PWK/PhJ. All mice were kept in temperature- and humidity-controlled rooms, in a 12/12 hr light/dark cycle, with lights on from 7:00 AM to 7:00 PM. Food and water were provided *ad libitum*. All procedures were approved by the IACUC at Yale University School of Medicine.

### 4.2 Audio Acquisition

Mice were placed inside a box (40 x 40 x 40 cm) with fresh bedding and covered by anechoic material (2” Wedge Acoustic Foam, Auralex) in order to attenuate external noise. Four boxes were recorded simultaneously, each one containing one mouse. Audio files were recorded using the recorder module UltraSoundGate 416H and a condenser ultrasound microphone CM16/CMPA (Avisoft Bioacoustics, Berlin, Germany) placed 15 cm above the animal. The experiments were recorded with a sampling rate of 250 kHz. The recording system had a flat response for sounds within frequencies between 20 kHz and 140 kHz, preventing distortions for the frequency of interest. The recordings were made by using Avisoft RECORDER 4.2 (version 4.2.16; Avisoft Bioacoustics) in a Laptop with an Intel i5 2.4 GHz processor and 4 GB of RAM. Using these settings, ten minutes of audio recording generated files of approximately 200 MB.

### 4.3 Spectral power

USVs were segmented on the audio files by analysis of their spectrograms. Aiming the configuration that would grant us the best time-frequency resolution, the spectrograms were calculated through a short-time Fourier transformation (STFT) using the following parameters: 1024 sampling points to calculate the discrete Fourier transform (NFFT = 1024),

Hamming window with length 256 and half-overlapping with adjacent windows to reduce artifacts at the boundary. The mathematical expression that gives us the short-time Fourier Transform is shown below:

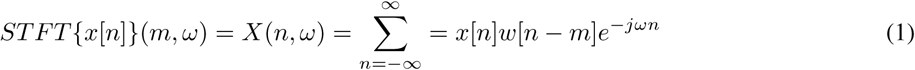

where the original signal *x*[*n*] is divided in chunks by the windowing function *w*[*m*]. The Fourier Transformation of the chunks result in a matrix with magnitude and phase for each time-frequency point.

The spectral power density, represented in the logarithmic unit decibels, is then given by

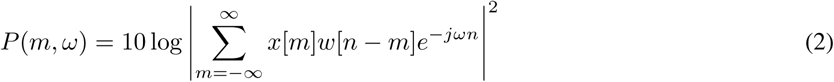

We used a high pass filter (45 kHz) to eliminate sources of noise in the audible range and to reduce the amount of data stored [16].

### 4.4 Normalization and contrast enhancement

Since USVs present higher intensity than the background and to avoid setting a fixed threshold for USV segmentation, we used contrast adjustment to highlight putative USV candidates and to reduce the variability across audio files. Contrast adjustment was obtained according to the following re-scaling equation:

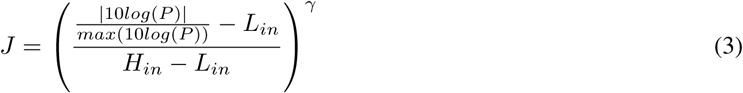

where *H*_*in*_ and *L*_*in*_ are the highest and the lowest intensity values of the adjusted image, respectively, and *P* is the power spectrum for each time-frequency point (pixel of the image). The parameter *γ* describes the shape of the mapping function between the original and the corrected image, such that *γ <* 1 results in darker pixels and *γ >* 1 in brighter pixels. We used a linear mapping for our application (*γ* = 1).

### 4.5 Adaptive thresholding and morphological operations

Due to non-stationary background noise and dynamic changes in the intensity of USVs within and between the audio files, we use adaptive thresholding methods to binarize the spectrograms. The threshold is computed for each pixel using the local mean intensity around the neighborhood of the pixel [4]. This method preserves hard contrast lines and ignores soft gradient changes. The integral image consists of a matrix *I*(*x, y*) that stores the sum of all pixel intensities *f* (*x, y*) to the left and above the pixel (*x, y*). The computation is given by the following equation:

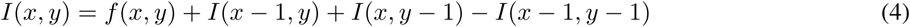

Therefore, the sum of the pixels values for any rectangle defined by a lower right corner (*x*_2_, *y*_2_) and upper left corner (*x*_1_, *y*_1_) is given as:

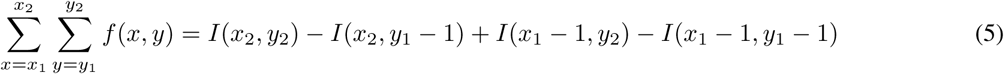

Then, the method computes the average of an *s × s* window of pixels centered around each pixel. The average is calculated considering neighboring pixels on all sides for each pixel. If the value of the current pixel is *t* percent less than this average, then it is set to black; otherwise it is set to white, as shown in the following equation:

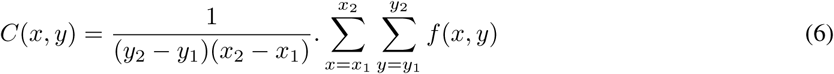

where *C*(*x, y*) represents the average around the pixel (*x, y*).

The binarized image is then constructed such as that pixels (*x, y*) with intensity *t* percent lower than *C*(*x, y*) are set to black [4]:

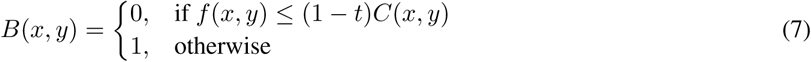

where *t* represents the sensitivity factor, and it was empirically chosen as *t* = 0.2 for our application. The segments are then subjected to a sequence of morphological operations: (i) opening (erosion followed by a dilation) with a rectangle 4 × 2 pixels as kernel; (ii) dilation with a line of length *l* = 4 and ∠ 90^*°*^ relative to the horizontal axis as kernel; (iii) filtering out candidates (i.e., dense set of white pixels) with < 60 pixels (correspondent to approximately 2 ms syllable); and (iv) dilation with a line of length *l* = 4 and ∠ 0^*°*^, making the USV candidates proportional to their original shape.

### 4.6 Local Median Filter

Noise due to the segmentation process is in the form of pixels or aggregate of pixels that are not associated with an event in the recording (a real USV or external noise) and are part of the pool of USV candidates. To determine if a USV candidate is relevant for further analysis, we perform a test - Local Median Filter - to compare the median intensity of the pixels in the USV candidate *k* (referred to as 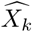) to the intensity of the pixels in a window that contains the candidate (referred to as 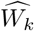). The Local Median Filter then determines if a USV candidate *k* is discarded based on the cumulative distribution of intensity ratio over all the USV candidates detected in the audio file 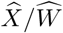. The bounding box that defines the window *W*_*k*_ is a rectangle with its four vertices defined as a function of the frequencies (*F*_*k*_) for USV candidate *k* and its time stamps (*T*_*k*_). Thus, the bounding box is defined as follows:

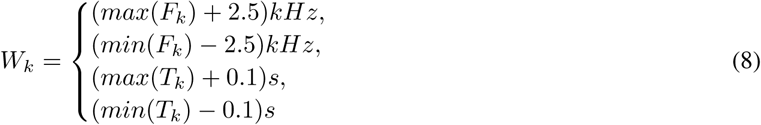

As seen in Equation 8, a 200 ms interval is analyzed around the USV candidate. Such a wide interval may present more than one USV in *W*_*k*_. However, the amount of pixels in *X*_*k*_ represents only 2.43 ± 0.10 % (mean ± SEM; median = 1.27, 95% CI [2.22, 2.63]; n = 59,781 images analyzed) of the total number of pixels contained in the window *W*_*k*_. Given this proportion between the number of pixels in *X*_*k*_ and *W*_*k*_, the median of the intensity distribution of the whole window 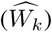 tends to converge to the median intensity of the background.

We used the ratio 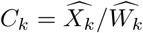 to exclude USV candidates that correspond to segmentation noise. We first calculated the cumulative distribution function (CDF) of *C*_*k*_ over all the USV candidates in an audio file (now referred to as ϒ). To find the inflection point in ϒ, a second-order polynomial fit for every set of 3 consecutive points was used to obtain local parametric equations (ϒ(*t*) = (*x*(*t*), *y*(*t*))) describing the segments of ϒ. Since the calculation of the inflection point is done numerically, the number of points chosen for this calculation should be such that we can have as many points of curvature as possible while preserving information of local curvature. Then, after a screening for the best number of points, ϒ was down-sampled to 35 equally spaced points and the inflection point was calculated. Using the local parametric equations, we calculated the tangent and normal vectors on each of the 35 points. Using these vectors, we estimated the changing rate of the tangent towards the normal at each point, which is the curvature *κ* [27] and can be calculated as follows:

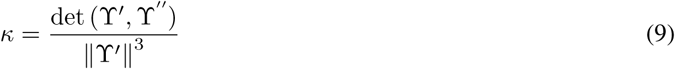

or by using the parametric equations:

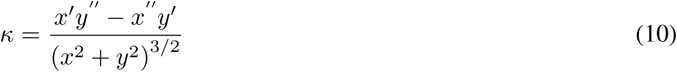

The inflection point is then determined as the point with maximum curvature of the CDF curve, and adopted as threshold *τ*. This threshold is calculated individually for each audio file since it can vary according to the microphone gain and the distance of the microphone from the sound source. In audio files with a very low number of USVs, the point of maximum curvature of the CDF curve was not detected, and no *τ* was estimated. In these cases, a default threshold *τ* = 0.92 was adopted as a conservative threshold, since no audio file presented inflection point as high as 0.92 in our *training* set. Only the USV candidates satisfying Equation 11 are kept for further analysis.

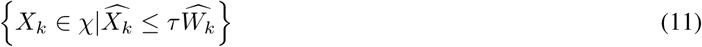

where *χ* represents the set of USV candidates that survived the Local Median Filter. Of note, the intensity of each pixel is calculated in decibels, which is given in negative units due to the low power spectrum.

### 4.7 Convolutional Neural Networks for USV classification

We use Convolutional Neural Networks to eliminate external noise from the pool of USV candidates and classify USVs in distinct types (see below). We use a transfer learning approach with an AlexNet [20] model pre-trained on the ImageNet dataset, and perform end-to-end training using our USV datasets. Briefly, the last three layers of the network were replaced in order to handle a twelve-categories classification task for our dataset (eleven *USV types* + *noise*).

The outputs of the segmentation process with detected USV candidates were centralized in windows of 220 ms. These windows were twice the maximum duration of USVs observed in mice [16] and were framed in individual 227 × 227 pixels images. Each image was then manually labeled by an experienced experimenter as noise (including acoustic or segmentation noise) or one of the USV categories. The labeled dataset was used to train the CNN to classify the USV candidates.

The images in our dataset were manually labeled according to our definitions of USV classes (adapted from [31] and [16]). The USV classes are described below:

**Complex:** 1-note syllables with two or more directional changes in frequency > 6 kHz. A total of 350 images were used for training.

**Step up:** 2-notes syllables in which the second element was ≥ 6 kHz higher from the preceding element and there was no more than 10 ms between steps. A total of 1,814 images were used for training.

**Step down:** 2-notes syllables in which the second element was ≥ 6 kHz lower from the preceding element and there was no more than 10 ms between steps. A total of 389 images were used for training.

**Two steps:** 3-notes syllables, in which the second element was ≥ 6 kHz or more different from the first, the third element was ≥ 6 kHz or more different from the second and there was no more than 10 ms between elements. A total of 701 images were used for training.

**Multiple steps:** 4-notes syllables or more, in which each element was ≥ kHz or more different from the previous one and there was no more than 10 ms between elements. A total of 74 images were used for training.

**Up-frequency modulation:** Upwardly frequency modulated with a frequency change ≥ 6 kHz. A total of 1,191 images were used for training.

**Down-frequency modulation:** Downwardly frequency modulated with a frequency change ≥ 6 kHz. A total of 1,775 images were used for training.

**Flat:** Constant frequency syllables with modulation ≤ 5 kHz and duration ≥ 12 ms. A total of 1,134 images were used for training.

**Short:** Constant frequency syllables with modulation ≤ 5 kHz and duration ≤ 12 ms. A total of 1,713 images were used for training.

**Chevron:** Shaped like an inverted *U* in which the peak frequency was ≥ 6 kHz than the starting and ending frequencies. A total of 1,594 images were used for training.

**Reverse chevron:** Shaped like an *U* in which the peak frequency was ≥ 6 kHz than the starting and ending frequencies. A total of 136 images were used for training.

**Noise:** Any sort of mechanical or segmentation noise detected during the segmentation process as a USV candidate. A total of 2,083 images were used for training.

In order to purposely create some overlap between the categories, USV with segments oscillating between 5 and 6 kHz were not defined or used for training. The assumption is that the CNN should find its transition method between two overlapping categories.

Our *training* dataset consisted of 12,954 images, wherein 2,083 were labeled as noise. This dataset correspond to mice of different strains (C57Bl6/J, NZO/HlLtJ, 129S1/SvImJ, NOD/ShiLtJ, and PWK/PhJ) and ages (5, 10, and 15 days of age) from both genders.

The CNN was trained using stochastic gradient descent with momentum, a batch size of M=128 images, and with a maximum number of epochs set to 100. Through a screening process for the set of hyper-parameters that would maximize the average performance of the network, the chosen learning rate was *α* = 10^−4^, momentum of 0.9, and weight decay *λ* = 10^−4^. To validate the training performance, each dataset was split into two disjoint sets; *training* set (90%) and a *validation* set (10%). The training and validation sets were independently shuffled at every epoch during training. The training was set to stop when the classification accuracy on the validation set did not improve for 3 consecutive epochs. When running in a GeForce GTX 980 TI, the final validation accuracy was 95.28% after 17 minutes of training.

### 4.8 Testing detection performance

To evaluate the performance of VocalMat, neonatal mice were recorded for 10 minutes upon social isolation in different conditions (Table 1) to increase the variability of the data. The spectrograms were manually inspected for the occurrence of USVs. The starting time for the detected USVs was recorded. USVs automatically detected by VocalMat with a start time matching manual annotation (±5 ms of tolerance) were considered correctly detected. USVs manually detected with no correspondent USV given by VocalMat were considered *false negative*. The false negatives were originated from missed USVs or USVs that the software labeled as noise. Finally, USVs registered by VocalMat without a correspondent in the manual annotation were considered *false positive* (see Table 2). In order to compare VocalMat to the other tools available, the same metrics were applied to the output of Ax [22], MUPET [36]. USVSEG [34], and DeepSqueak [10].

**Table 2:** Summary of possible outcomes for the detection validation

| Manual | Automated | Actual meaning | Label |
| --- | --- | --- | --- |
| Detected | Detected | Success | True positive |
| Detected | Not detected | Missed or classified as noise | False negative |
| Not detected | Detected | Noise | False positive |

### 4.9 Diffusion maps for output visualization

One of the main characteristics of VocalMat is the possibility of classifying USVs as a distribution of probabilities over all the possible labels. Since we classify USV candidates in 11 categories, to have access to the distribution of probabilities, we would need to visualize the data in 11 dimensions. Here, as an example of analytical methods that can be applied to the output data from VocalMat, we used Diffusion Maps [11] to reduce the dimensionality of the data to three dimensions. Diffusion Maps allows remapping of the data into a Euclidean space, which ultimately results in a clustering of USVs based on the similarity of their probability distribution. A Gaussian kernel function defines the connectivity between two data points in a Euclidean manifold. Such kernel provides the similarity value between two data points *i* and *j* as follows:

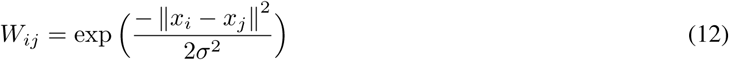

where *W*_*ij*_ represents the similarity value between observations *i* and *j*. The parameter *σ* corresponds to the bandwidth, and it is set based on the average Euclidean distance observed between observations of the same label. For our application, *σ* = 0.5 was set based on the distance distribution observed in our data.

The similarity matrix is then turned into a probability matrix by normalizing the rows:

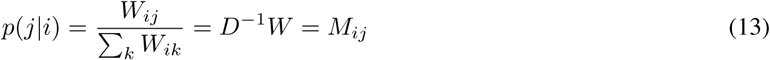

where ∑_*k*_ *W*_*ik*_ = *D*_*ii*_ has the row sum of *W* along its diagonal. The matrix *M* gives the probability of walking from node *i* to any other node. In other words, the probability that the USV *i* is close to another USV *j* given their probability distribution.

Once we take one step in such Euclidean space, the probabilities are updated, since the set of likely nodes for the next move are now updated. This idea of moving from node to node while updating the probabilities gives us a “diffused map”.

The process of moving from a USV *i* to *j* after *t* steps in this Euclidean space is computed as follows:

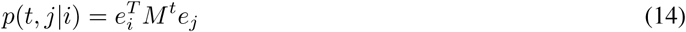

For our application, we use *t* = 2.

Next, we find the coordinate functions to embed the data in a lower-dimensional space. The eigenvectors of *M* give such a result. Because *M* is not symmetric, the eigendecomposition is computed through SVD decomposition [15]:

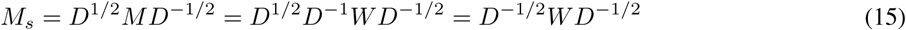

and since *D*^−1*/*2^ and *W* are symmetric, *M*_*s*_ is also symmetric and allows us to calculate its eigenvectors and eigenvalues. For the sake of notation, consider:

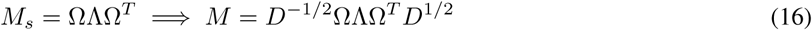

Considering Ψ = *D*^−1*/*2^Ω (right eigenvectors of *M*) and Φ = *D*^1*/*2^Ω (left eigenvectors of *M*), we verify that Φ^*T*^ = Ψ^−1^, therefore they are mutually orthogonal and *M* and *M*_*s*_ are similar matrices. Thus,

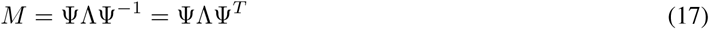

and the diffusion component shown in Equation 14 is incorporated as the power of the diagonal matrix composed by the eigenvalues of *M* :

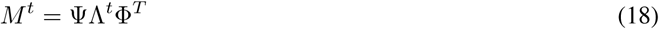

We use the scaled right eigenvectors by their corresponding eigenvalues (Γ = ΨΛ) as the coordinate functions. Since the first column of Γ is constant across all the observations, we use the 2nd to 4th coordinates in our work.

### 4.10 Vocal repertoire analysis via manifold alignment

The result of the embedding by Diffusion Maps allows 3D visualization of the probability distribution for the USVs. The direct comparison of different 3D maps is challenging to obtain as the manifolds depend on data distribution, which contains high variability in experimental samples. To address this problem and compare the topology of different manifolds, we considered this a transfer learning problem [28]. We used a manifold alignment method for heterogeneous domain adaptation [37, 35]. Using this method, two different domains are mapped to a new latent space, where samples with the same label are matched while preserving the topology of each domain.

We used the probability distribution for the USVs for each dataset to build the manifolds [37]. Each manifold was represented as a Laplacian matrix constructed from a graph that defines the connectivity between the samples in the manifold. The Laplacian matrix is then defined as *L* = *W*_*ij*_ − *D*_*ii*_ (see Equation 12).

The final goal is to remap all the domains to a new shared space such that samples with similar labels become closer in this new space. In contrast, samples with different labels are pushed away while preserving the geometry of the manifolds. It leads to the necessity of three different graph Laplacians: *L*_*s*_ (relative to the similarity matrix and responsible for connecting the samples with the same label), *L*_*d*_ (dissimilarity matrix and responsible for connecting the samples with different labels), and *L* (similarity matrix responsible for preserving the topology of each domain). [37] show that the embedding that minimizes the joint function defined by the similarity and dissimilarity matrices is given by the eigenvectors corresponding to the smallest non-zero eigenvalues of the following eigendecomposition:

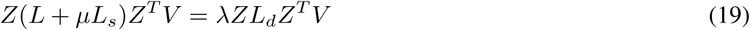

where *Z* is a block diagonal containing the data matrices 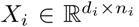, (*n*_*i*_ samples and *d*_*i*_ dimensions for the *i*^*th*^ domain) from the two domains. Thus, *Z* = *diag*(*X*_1_, *X*_2_). The matrix *V* contains the eigenvectors organized in rows for each domain, *V* = [*v*_1_, *v*_2_]^*T*^. The *µ* is weight parameter, which goes from preserving both topology and instance matching equally (*µ* = 1) or focusing more on topology preservation (*µ >* 1).

From Equation 19, we then extract 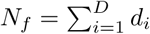 features, and the projection of the data to this new common space *ℱ* will be given by

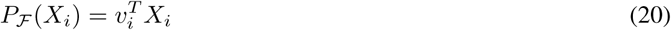

To measure the performance of the alignment, linear discriminant analysis (LDA) [21] is used to show the ability to project the domains in a joint space. The LDA is trained on half of the samples in order to predict the other half. The error of the alignment is given as the percentage of samples that would be misclassified when projected into the new space (overall accuracy) [35].

Another measurement to quantify the quality of the alignment is by calculating the agreement between the projections, which is given by Cohen’s Kappa coefficient (*κ*) [1]. In this method, the labels are treated as categorical, and the coefficient compares the agreement with that expected if ratings were independent. Thus, disagreements for labels that are close are treated the same as labels that are far apart.

Cohen’s coefficient is defined as:

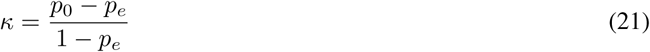

where *p*_0_ is the observed agreement (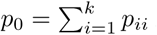 for a confusion matrix *p* = *n/N*, in which *n* is the raw confusion matrix and *N* is the total number of samples, composed by the projection of the *k* labels), which corresponds to the accuracy; *p*_*e*_ is the probability of agreement by chance (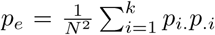, where *p*_*i*_ is the number of times an entity of label *i* was labeled as any category and *p*_.*i*_ is the number of times any category was predicted as label *i*). Therefore, a *κ* = 0 represents no agreement (or total misalignment of manifolds) and *κ* = 1 is a total agreement.

In this context, the overall accuracy (*OA*) is given by 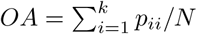, where *N* is the total number of samples.

The asymptotic variance for *κ* is given as follows:

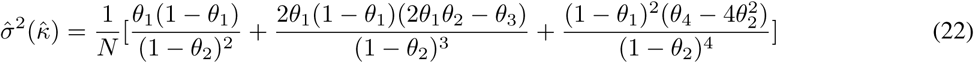

where

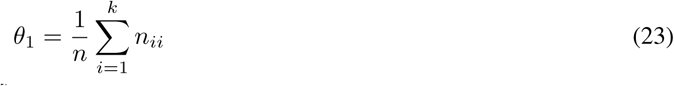

(which turns into accuracy once it is divided by *N*),

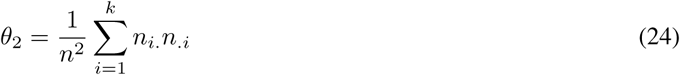

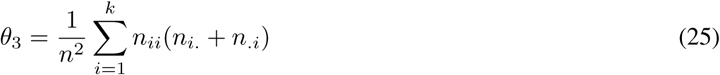

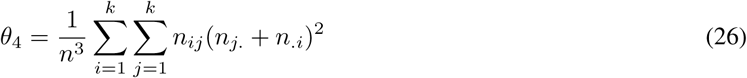

From Equation 22 we can calculate the Z-score, which can express the significance of our *κ*:

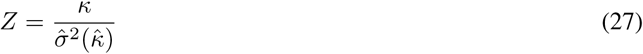

And the 95% confidence interval as

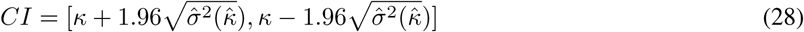

The third form of error measurement is the evaluation of the projection per USV class from each domain remapped into the new space. This method is based on the fact that this new space is the one in which the cost function expressed by Equation 19 is minimized and, therefore, the projection from each domain into the new space has its projection error for each class. As a consequence, the mean of the projection error from each domain to the new space for each class can be used as a quantitative measurement of misalignment of projected domains.

### 4.11 Quantification and statistical analysis

MATLAB (2019a or above) and Prism 8.0 were used to analyze data and plot figures. All figures were edited in Adobe Illustrator CS6/CC. Data were first subjected to a normality test using the D’Agostino & Pearson normality test or the Shapiro-Wilk normality test. When homogeneity was assumed, a parametric analysis of variance test was used. The Student’s t test was used to compare two groups. The Mann-Whitney U test was used to determine significance between groups. Two sample Kolmogorov–Smirnov test was used to calculate the statistical differences between the contrast of USVs and noise. Statistical data are provided in text and in the figures. In the text, values are provided as mean ± SEM. p < 0.05 was considered statistically significant. The 95% confidence intervals are reported in reference to the mean. The true positive rate is computed as the ratio between true positive (hit) and real positive cases. The true negative rate is the ratio between true negative (correct rejection) and real negative cases. The false negative rate is the ratio between false negative (type I error) and real positives cases. The false positive (type II error) is the ratio between false positive and real negative cases. The false discovery rate is the ratio between false positive and the sum of false positives and real positives.

### 4.12 Code and data availability

VocalMat is available on GitHub (https://github.com/ahof1704/VocalMat.git) for academic use. Our dataset of vocalization images is available on OSF (https://osf.io/bk2uj/).

## 5 Acknowledgments

We thank lab members for critical data collection and insights in the manuscript. M.O.D. was supported by a NARSAD Young Investigator Grant ID 22709 from the Brain & Behavior Research Foundation, by the National Institute Of Diabetes And Digestive And Kidney Diseases of the National Institutes of Health (R01DK107916), by a pilot grant from the Yale Diabetes Research Center (P30 DK045735), by the Yale Center for Clinical Investigation Scholar Award, by the Whitehall Foundation, by the Charles H. Hood Foundation, Inc. (Boston, MA), by a pilot grant from the Modern Diet and Physiology Research Center (The John B. Pierce Laboratory), by a grant of the Foundation for Prader-Willi Research, and by the Reginald and Michiko Spector Award in Neuroscience. M.O.D. also received support from the Conselho Nacional de Desenvolvimento Científico e Tecnológico (CNPq) and Coordenadoria de Aperfeiçoamento de Pessoal de Nível Superior (CAPES), Brazil. A.F. and G.M.S. were partially supported by scholarships from CAPES during the development of this project. The authors declare no conflict of interest.

## 6 Supplementary material

**Figure S1:**
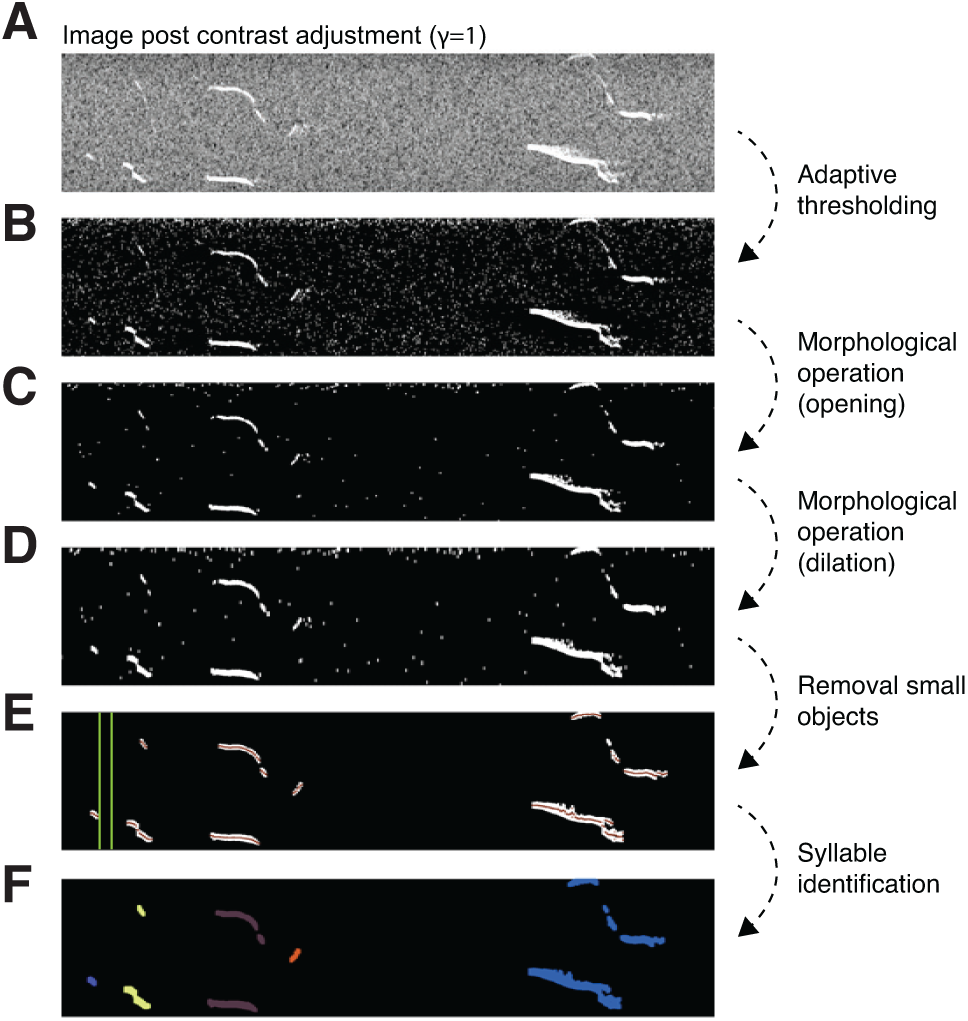
Image processing pipeline for segmentation of USVs in spectrograms. **(A)** Segment of a spectrogram post contrast adjustment (*γ* = 1). **(B)** Output image post binarization using adaptive thresholding. **(C)** Resulting image from the opening operation with rectangle 4×2. **(D)** Result from the dilation with line *l*=4 and ∠ 90^*°*^. **(E)** Removal of too small objects (≤ 60 pixels), mean of cloud points for each detected USV candidate being shown in red and green lines shows an interval of 10 ms. **(F)** Result after separating syllables based on the criterion of maximum interval between two tones in a syllable. The different colors differentiate the syllables from each other.

**Table S1:**
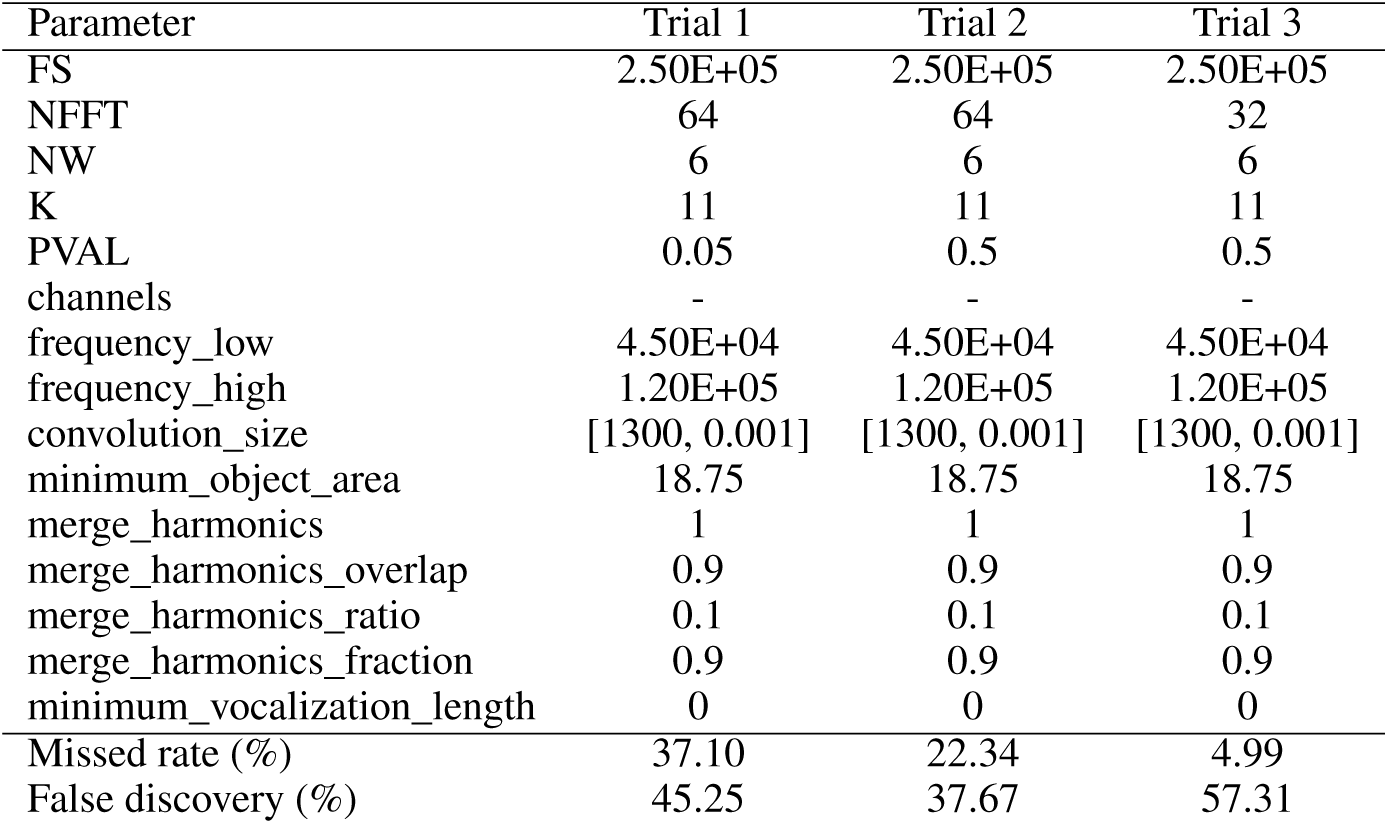
List of parameters and performance for Ax

| Parameter | Trial 1 | Trial 2 | Trial 3 |
| --- | --- | --- | --- |
| FS | 2.50E+05 | 2.50E+05 | 2.50E+05 |
| NFFT | 64 | 64 | 32 |
| NW | 6 | 6 | 6 |
| K | 11 | 11 | 11 |
| PVAL | 0.05 | 0.5 | 0.5 |
| channels | - | - | - |
| frequency_low | 4.50E+04 | 4.50E+04 | 4.50E+04 |
| frequency_high | 1.20E+05 | 1.20E+05 | 1.20E+05 |
| convolution_size | [1300, 0.001] | [1300, 0.001] | [1300, 0.001] |
| minimum_object_area | 18.75 | 18.75 | 18.75 |
| merge_harmonics | 1 | 1 | 1 |
| merge_harmonics_overlap | 0.9 | 0.9 | 0.9 |
| merge_harmonics_ratio | 0.1 | 0.1 | 0.1 |
| merge_harmonics_fraction | 0.9 | 0.9 | 0.9 |
| minimum_vocalization_length | 0 | 0 | 0 |
| Missed rate (%) | 37.10 | 22.34 | 4.99 |
| False discovery (%) | 45.25 | 37.67 | 57.31 |

**Figure S2:**
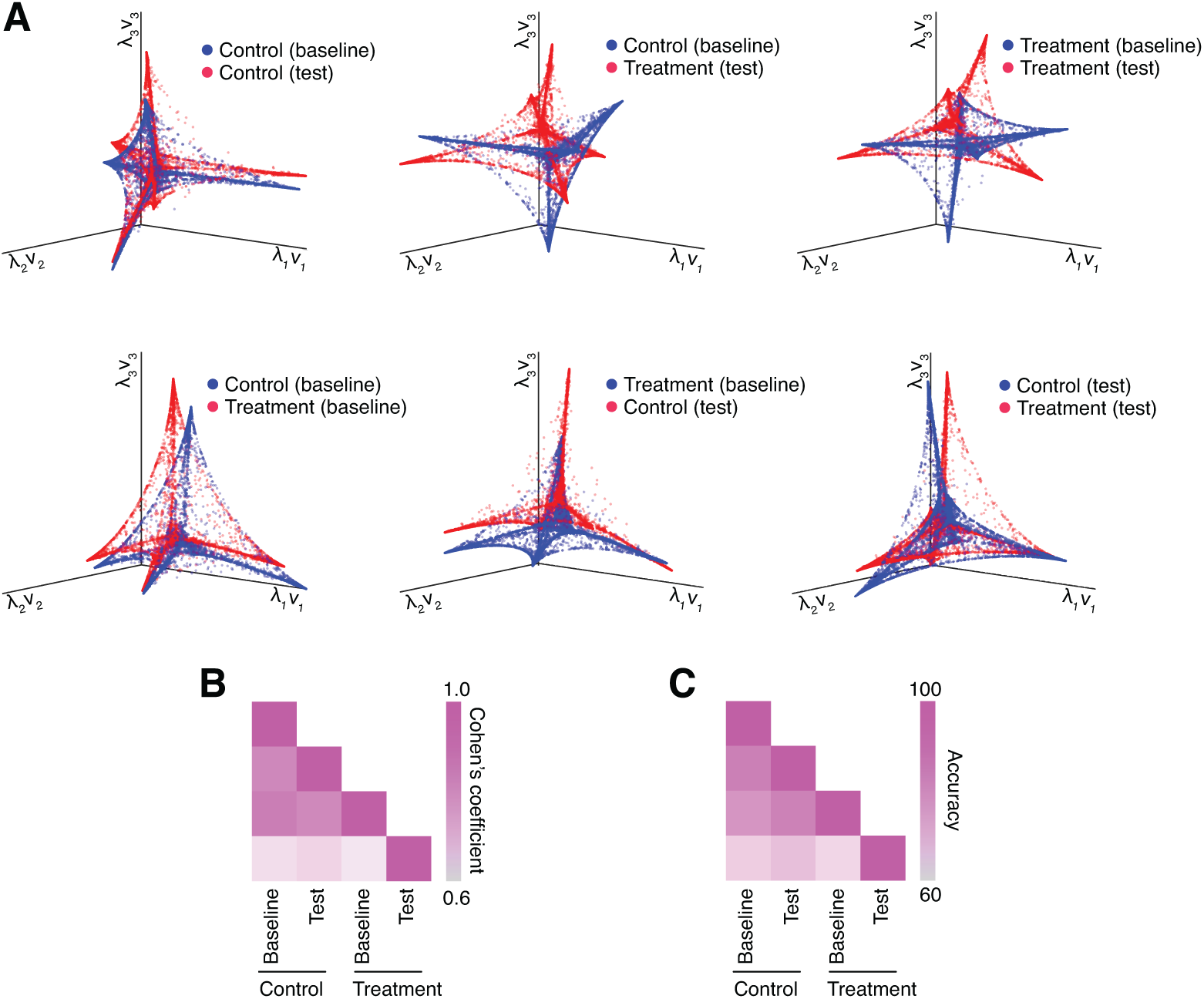
Alignment of the manifolds between pairs of experimental conditions. (**A**) Illustration of the resulting manifold alignment for each pair of experimental conditions. The quality of the alignment between the manifolds is assessed by (**B**) Cohen’s coefficient and (**C**) overall projection accuracy into joint space.

**Table S2:** List of parameters and performance for MUPET

| Parameter | Trial 1 | Trial 2 | Trial 3 | Trial 4 | Trial 5 | Trial 6 | Trial 7 | Trial 8 |
| --- | --- | --- | --- | --- | --- | --- | --- | --- |
| noise-reduction | 5 | 5 | 5 | 1 | 1 | 0.5 | 2 | 1 |
| min-syllable-duration | 2 | 2 | 2 | 2 | 2 | 2 | 2 | 2 |
| max-syllable-duration | 200 | 200 | 200 | 200 | 200 | 200 | 200 | 200 |
| min-syllable-total-energy | -15 | -15 | -25 | -25 | -10 | -25 | -25 | -35 |
| min-syllable-peak-amplitude | -25 | -25 | -35 | -35 | -16 | -35 | -35 | -45 |
| min-syllable-distance | 5 | 10 | 10 | 10 | 10 | 10 | 10 | 10 |
| Missed rate (%) | 41.84 | 44.92 | 44.19 | 34.63 | 41.05 | 33.74 | 37.72 | 34.63 |
| False discovery (%) | 38.78 | 40.02 | 41.92 | 52.74 | 51.08 | 53.11 | 51.07 | 53.07 |

**Table S3:**
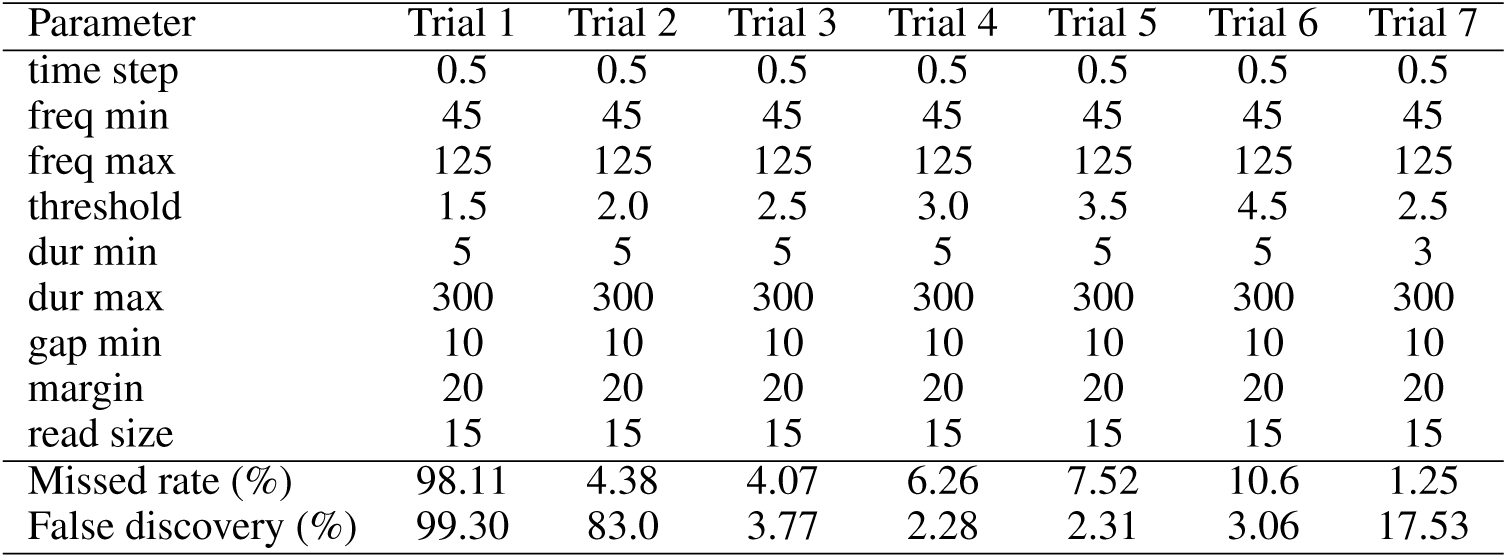
List of parameters and performance for USVSEG

**Table S4:** List of parameters and performance for DeepSqueak

| Parameter | Value |
| --- | --- |
| overlap | 0.1 |
| frequency cut off high | 120 |
| frequency cut off low | 45 |
| neural network | MouseCall_Network_V2 |
| detection | normal |
| Missed rate (%) | 27.13 |
| False discovery (%) | 7.61 |

**Table S5:** VocalMat accuracy per class

| Type | N | Mean ± SEM (%) | Median [95% CI] (%) |
| --- | --- | --- | --- |
| Step up | 902 | 83.58 ± 6.50 | 91.56 [66.85, 100.00] |
| Chevron | 758 | 85.37 ± 3.93 | 85.28 [75.25, 85.48] |
| Two steps | 579 | 74.41 ± 4.16 | 70.47 [63.71, 85.11] |
| Down-FM | 557 | 90.74 ± 1.23 | 90.83 [87.56, 93.91] |
| Up-FM | 485 | 88.04 ± 2.38 | 87.59 [81.90, 94.17] |
| Short | 358 | 88.28 ± 1.88 | 89.62 [83.45, 93.11] |
| Complex | 281 | 76.64 ± 3.72 | 76.24 [67.07, 86.22] |
| Flat | 190 | 84.20 ± 4.14 | 83.51 [73.56, 94.84] |
| Step down | 142 | 84.74 ± 4.60 | 83.77 [72.90, 96.58] |
| Mult. steps | 80 | 45.89 ± 10.70 | 38.10 [16.18, 75.61] |
| Rev. Chevron | 61 | 65.18 ± 14.17 | 73.87 [28.74, 100.00] |
| Noise | 511 | 96.67 ± 0.55 | 96.67 [95.23, 98.10] |

**Table S6:** VocalMat accuracy considering the two most likely labels

| Type | N | Mean ± SEM (%) | Median [95% CI] (%) |
| --- | --- | --- | --- |
| Step up | 902 | 91.64 ± 4.86 | 97.18 [79.15, 100.00] |
| Chevron | 758 | 96.08 ± 1.36 | 97.20 [92.57, 99.58] |
| Two steps | 579 | 91.43 ± 1.80 | 91.77 [86.79, 96.06] |
| Down-FM | 557 | 97.08 ± 0.98 | 96.93 [94.57, 99.59] |
| Up-FM | 485 | 96.25 ± 1.40 | 97.30 [92.66, 99.84] |
| Short | 358 | 96.53 ± 1.12 | 96.72 [93.66, 99.41] |
| Complex | 281 | 92.10 ± 2.30 | 91.44 [86.20, 98.00] |
| Flat | 190 | 94.21 ± 3.96 | 97.73 [84.02, 100.00] |
| Step down | 142 | 96.11 ± 1.99 | 97.96 [91.01, 100.00] |
| Mult. steps | 80 | 83.64 ± 7.33 | 85.71 [63.28, 100.00] |
| Rev. Chevron | 61 | 77.65 ± 15.75 | 91.29 [37.17, 100.00] |
| Noise | 511 | 98.00 ± 0.45 | 97.87 [96.84, 99.17] |

